# Accurate flux predictions using tissue-specific gene expression in plant metabolic modeling

**DOI:** 10.1101/2022.09.05.506655

**Authors:** Joshua A.M. Kaste, Yair Shachar-Hill

## Abstract

**Motivation:** The accurate prediction of complex phenotypes such as metabolic fluxes in living systems is a grand challenge for systems biology and central to efficiently identifying biotechnological interventions that can address pressing industrial needs. The application of gene expression data to improve the accuracy of metabolic flux predictions using mechanistic modeling methods such as Flux Balance Analysis (FBA) has not been previously demonstrated in multi-tissue systems, despite their biotechnological importance. We hypothesized that a method for generating metabolic flux predictions informed by relative expression levels between tissues would improve prediction accuracy.

**Results:** Relative gene expression levels derived from multiple transcriptomic and proteomic datasets were integrated into Flux Balance Analysis predictions of a multi-tissue, diel model of Arabidopsis thaliana’s central metabolism. This integration dramatically improved the agreement of flux predictions with experimentally based flux maps from 13C Metabolic Flux Analysis (MFA) compared with a standard parsimonious FBA approach. Disagreement between FBA predictions and MFA flux maps, as measured by weighted averaged percent error values, dropped from between 169-180% and 94-103% in high light and low light conditions, respectively, to between 10-12% and 9-11%, depending on the gene expression dataset used. The incorporation of gene expression data into the modeling process also substantially altered the predicted carbon and energy economy of the plant.

**Availability:** Code is available from https://github.com/Gibberella/ArabidopsisGeneExpressionWeights

## Introduction

A grand challenge for systems biology is the ability to accurately predict complex phenotypes from omic datasets based on functional principles and mechanisms. Patterns of cellular metabolism – flux maps – are one such complex phenotype (1), for which tools to predict phenotypes from basic assumptions have proven useful in exploring and designing metabolic capabilities (2–4). Methods to quantify flux maps from labeling data now allow the testing of such predictions in both simpler and multicellular systems. However, the integration of omic data to improve the accuracy of flux predictions is still at an early stage.

Metabolic flux predictions are also important for real world applications since modifying an organism’s metabolic activity in order to achieve some practical aim, such as overproducing a specific metabolite, is central to many biotechnology projects. As in other areas of engineering, metabolic engineering can benefit from mathematical models that describe and predict the behavior of the relevant system(s). Researchers have developed two major modeling approaches to address this need: (i) 13C-Metabolic Flux Analysis (13C-MFA) and (ii) Flux Balance Analysis (FBA) (2, 5). With 13C-MFA, steady-state or kinetic isotopic labeling data for metabolites in a small-to medium-sized network are used to obtain estimates of the net and exchange fluxes through that network (5). These metabolic flux maps are regarded as the most reliable measures of *in vivo* metabolic fluxes; however, the throughput of this technique is limited by the large amounts of isotopic labeling data and other measurements needed to generate each flux map. FBA, which is based on applying conservation principles to a network of reactions using one or more assumptions about the functional objective(s) driving biological organization, requires substantially less experimental input data, and is therefore an attractive and commonly used metabolic modeling technique.

FBA and related metabolic modeling methods in microbial systems, together with Genome-Scale Metabolic Models (GEMs) that represent the biochemical reactions encoded in an organism’s genome, have enabled radical modification of microbial central metabolism (e.g. Gleizer *et al*., 2019) and substantial improvements in bioproduct yields (e.g. Jin *et al*., 2007; Lee *et al*., 2007) These methods can, for example, allow bioengineers to predict the behavior of their system and identify interventions, such as gene knock-outs or knock-ins, that will help them modify the organism’s phenotype (4, 9). However, many metabolic engineering applications require the modification not of microorganisms, but of multicellular eukaryotes like plants. Most GEMs of plants to date (e.g. (10–13)), have treated plants, which are composed of multiple tissues with substantial functional diversity, as single-tissue aggregated metabolic networks. This has motivated the creation of “multi-tissue” GEMs to investigate source-sink dynamics and resource allocation, with the earliest efforts in this space focusing on the interplay between mesophyll and bundle-sheath cells in C4 photosynthesis (14, 15).

Re-engineering of plant metabolism on the scale seen in microbial systems has not, to-date, been possible and predictive modeling has been neither validated in detail nor applied to successful plant metabolic engineering. This is in part a result of the ease and high throughput of microbial transformation relative to that of even model plants like *Arabidopsis thaliana*. In addition to the greater experimental difficulty, the metabolic modeling of these systems is also substantially harder. There is, consequently, a relative lack of MFA datasets with which to compare the predicted flux maps from FBA in plants. This contrasts with the availability of rich multi-omic datasets combining flux estimates with transcript and protein data for a number of different genotypes and growth conditions in systems like *E. coli* (16). The substantial challenge involved in generating 13C-MFA flux maps for plants makes improvement in the quality of plant FBA flux predictions an attractive path towards replicating the biotechnological successes seen in microbes.

An appealing approach to improving the quality of plant FBA predictions is the integration of additional network-wide data from transcriptomic and proteomic datasets. Gene expression data – particularly transcript data – is substantially easier to generate than 13C-MFA flux maps. Previous attempts at the integration of gene expression datasets into metabolic flux predictions have been reviewed elsewhere (17, 18). A substantial number of methods developed before 2014 were evaluated on the basis of their ability to improve upon parsimonious FBA (pFBA) (19) in terms of their predictions’ agreement with MFA-estimated fluxes in microorganisms and were found to not do so reliably (18). A key limitation of these studies was a lack of comparison of FBA-predictions against 13C-MFA derived flux estimates. This lack of comparison against 13C-MFA is shared by the plant FBA literature, in which we are aware of only a small number of evaluations under heterotrophic conditions in green algae (20), *Arabidopsis* cell cultures (21, 22), and *Brassica napus* embryos (23). Since then, several studies have developed algorithms benchmarked by their ability to make predictions in agreement with empirical flux maps derived from MFA studies (24, 25). These studies have focused on unicellular organisms (*Escherichia coli* and *Saccharomyces cerevisiae*) studied under different conditions or genetic alteration. Their applicability to FBA in more complex systems is limited by the large number of resource-intensive MFA datasets needed to calibrate them (24) or their need for a reference expression dataset paired with an assumed-correct flux map (25).

In the interest of advancing the accuracy of FBA in systems with multiple cell types, particularly in plants with their complex metabolic networks, we have developed a method that allows for the integration of tissue-atlas data from multi-tissue systems into the flux-minimization procedure employed in pFBA. This method incorporates evidence from gene expression datasets into FBA metabolic flux predictions by applying weights to individual reactions according to the relative transcript or protein expression of the gene(s) assigned to those reactions between different modeled tissues. The method described in this study is evaluated on the basis of its ability to produce predictions in accordance with MFA flux maps. We demonstrate substantial improvements in the agreement of our FBA predicted fluxes with flux estimates from a 13C-MFA study on *Arabidopsis thaliana* rosette leaf central metabolism (26). Finally, we show that multiple gene expression datasets, when used as inputs, all result in a similar improvement in agreement and that this result generalizes across multiple different MFA-estimated flux maps. We believe that this approach has particular potential for plant and animal systems for which there are only a limited number of well-established experimental flux maps.

## Methods

### Overview of approach

Our method makes two key assumptions: (1) First, metabolic flux maps that are predicted from parsimonious FBA (19), minimizing the sum total of flux through the network, are more likely to reflect real flux maps than ones not subject to this constraint, and (2) A reaction present in two tissues *A* and *B* catalyzed by an enzyme encoded by a gene that is highly expressed in *A* and poorly expressed in *B* is likely to carry higher flux in tissue *A*.

We incorporate assumption 1 by making the objective function of our FBA optimization the minimization of total flux, the same as pFBA (19). This is represented mathematically as finding the minimum value of the linear combination of all fluxes in the network, with each flux v_i_ multiplied by a corresponding coefficient c_i_:

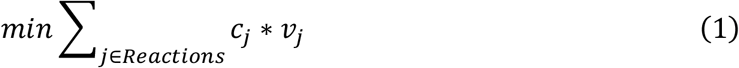

Where *Reactions* is the list of all reactions *j* in the network, *v*_*j*_ is the flux through a reaction *j*, and c_j_ is the coefficient – which we will hereafter refer to as a *penalty weight* since it represents a penalization on the likelihood of using a reaction *j* to carrying flux. When *c*_*j*_ takes a value of 1 for all reactions, our method reduces to pFBA, which can be seen as the limiting case of gene expression having no influence in predicting network flux patterns. We incorporate assumption 2 by calculating, for each reaction in our network model, a coefficient derived from the relative expression of genes encoding enzyme(s) that catalyze that reaction between the different tissues in our gene expression dataset. This use of the coefficient vector to account for relative expression evidence is similar to the approach taken in (27), but in this case relative expression is across tissues within a single multi-tissue model. The association between reactions and genes is captured by the Gene-Protein-Reaction (GPR) terms in the model. This results in reactions mapped to relatively highly expressed genes receiving small values of c_j_ and reactions mapped to minimally expressed genes receiving large ones.

### Multi-tissue diel model construction and dataset selection

The *Arabidopsis thaliana* core metabolism model developed in (12) was used as the basis for a multi-tissue diel model. This model was chosen due to its rich GPR annotation and focus on central metabolism. The core model was duplicated six times to create leaf, stem, and root versions of the model for both day and night, which were interconnected by transporters allowing the movement of specific compounds and metabolites. Additional details on the constraints applied to the model can be found in the *Supplementary Methods*. The full model used in this study can be found in **Supplemental Dataset 2**.

13C-MFA flux maps were obtained *in planta* in *Arabidopsis thaliana* by (26), and these were used as the empirical best estimates of flux distributions. The pairing of fluxes in the MFA network (26) to the FBA network are described in **Supplemental Dataset 1**.

We searched the literature for high-quality, high-coverage RNA-seq tissue atlases and quantitative proteomic tissue atlases and found two suitable datasets meeting these criteria: Merger *et al*. 2019 (28) and Klepikova *et al*. 2016 (29). For dataset IDs and bioinformatic processing details, see *Supplementary Methods*.

### Gene expression weight vector calculation

We calculated the penalty weight for each gene in each tissue on the basis of how the expression of a reaction in a particular tissue, as measured by transcriptomic or proteomic abundance, compared to the expression of that same gene in other modeled tissues.

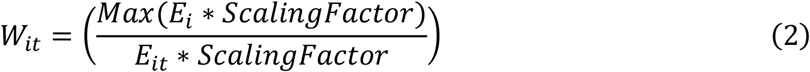

Where *W*_it_ is the weight for a given gene *i* in a tissue *t*, E_i_ is the list of expression values of gene *i* for each tissue, E_it_ is the expression of gene *i* in tissue *t*, Max() is the maximum value from a set of one or more elements, and the ScalingFactor is a coefficient that modulates the magnitude of the calculated weights. Many GPRs in the model consist of multiple genes that represent isozymes or members of protein complexes. The former are denoted by OR terms and the latter by AND terms in the GPR formulation. This results in many reactions having more than one penalty weight due to being mapped to multiple genes. We combine these multiple weights into a single value for each reaction by averaging the penalty weights of isozymes and taking the “worst” (i.e. largest, most penalizing value) when genes form subunits of a protein complex. As an example, the weight for a reaction *R* in the leaf subnetwork of our model with a GPR of the form (Gene1 OR Gene2) AND (Gene3), corresponding to a protein complex made of the product of Gene 3 and the product of either Gene 1 or Gene 2, would be represented by:

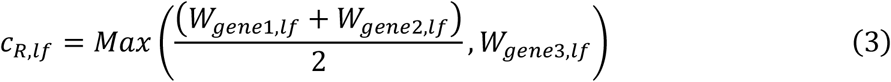

Where c_R,lf_ represents the overall weight for a reaction and W_gene1,lf_, W_gene2,lf_, and W_gene3,lf_ are the weights for the individual genes Gene1, Gene2, and Gene3. Note that in the present implementation of this method, stoichiometric coefficients in GPR terms are ignored. When the one or more genes contained in a GPR for a reaction/tissue combination are all more highly expressed than the same genes in the other tissues, the scale for that reaction/tissue combination will be 1. For reaction/tissue combinations that have no corresponding GPR, we explored setting the weights to 1 or a value calculated from the median weight assigned to reactions in the same tissue (for details, see *Supplementary Methods*).

### Optimization

The optimization done in this paper is a variation on pFBA, which finds the flux map(s) that satisfies imposed constraints with minimum total flux through the network (19). The minimization of total flux **(Eq. 1)** is subject to the following constraints:

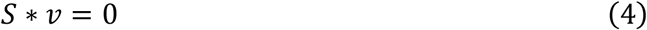

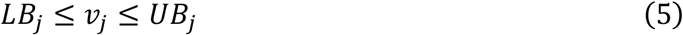

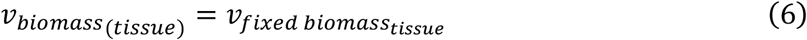

Where *S* is the stoichiometric matrix of the metabolic network being modeled, *v* is the vector of all fluxes, *LB* and *UB* are the vectors of all upper and lower bound constraints, and *v*_biomass(tissue)_ and *v*_fixed biomass(tissue)_ are the biomass flux for a given tissue and the defined biomass constraint for that tissue, respectively. Eq. 4 represents the steady state of all internal metabolites, Eq. 5 represents the imposition of bounds and reversibility constraints, and Eq. 6 represents the definition of biomass accumulation rates. All optimizations were done in the COnstraint-Based Reconstruction and Analysis (COBRA) Toolbox in MATLAB (30) using the Gurobi™ optimizer version 8.1.1 (31).

### Error evaluation

We make the assumption that the 13C-MFA fluxes reported in (26) are the true *in vivo* metabolic fluxes and therefore regard the discrepancy between FBA-predicted fluxes and these 13C-MFA fluxes as a measure of error. Biomass accumulation (i.e. the difference in dry weight between a timepoint t and another timepoint t_-1_) was not reported in (26), but is the basis for the flux through the biomass equation in FBA. To allow a comparison between our FBA-predicted fluxes and the MFA-estimated fluxes in (26), we set an arbitrary biomass flux of 0.01 g/hr through the leaf, stem, and root biomass reactions in both the day and night, similar to the approach taken in (32). We then normalized our fluxes by multiplying them by a factor A calculated as the ratio of the measured leaf CO_2_ uptake from (26) and the net leaf CO_2_ uptake in our FBA flux map. A weighted average error for each FBA-predicted flux map was then obtained using the following expression:

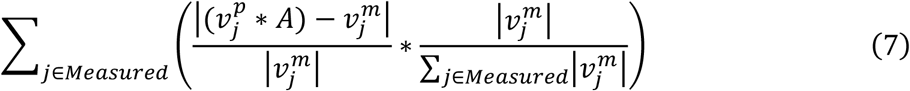

Where 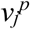 and 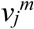 are the FBA-predicted and MFA-estimated fluxes of a flux *j* and A is the normalization factor previously described. We calculated weighted average errors rather than just average errors because small absolute differences between FBA-predicted and MFA-estimated flux values can correspond to extremely large % error values when the MFA-estimated fluxes are small. Additional details on the error evaluation done in this study can be found in the *Supplementary Methods*. We quantified the maximum/minimum weighted average errors of each flux map using Flux Variability Analysis (FVA) (33). For details, see *Supplementary Methods*.

## Results

### The application of gene expression weight reliably reduces discrepancies between FBA-predicted and MFA-estimated fluxes

Predicted flux maps were generated for a multi-tissue diel model of *Arabidopsis thaliana*’s central metabolism using flux balance analysis in which the sum of all the metabolic and transport fluxes required for steady state growth is minimized, with each flux being multiplied by a weight coefficient that was derived from the relative expression of the gene(s) involved in conducting that flux (see methods). Weights for each reaction were calculated from RNA-seq (28, 29) and proteomic (28) datasets using the relative expression of each gene in the different tissues. The weighted average % error between these flux maps and 13C-MFA estimates from (26) were used to quantify the accuracy of these FBA predictions, as compared to the accuracy of flux maps generated by pFBA (19) alone. The flux maps arrived at after the application of either transcriptomic or proteomic weights show greater agreement, as measured by the weighted average % error, with 13C-MFA estimates than the results from pFBA alone **(Table 1)**. These reductions in error are substantial and statistically significant at α = 0.01. These increases in agreement are consistent across comparisons against two different flux maps (high-light and low-light) and are sustained across a range of assumed ratios of starch to sucrose production and carboxylase to oxygenase fluxes through RuBisCO (vo/vc). Marked reductions in error are seen whether one uses the transcriptomic or proteomic tissue-atlas datasets from (28) or the transcriptomic dataset from (29), so that the improvement in flux predictions is not dependent on the values obtained in a specific gene expression dataset or type.

**Table 1.** Weighted average % error values calculated from weighted vs. unweighted flux maps for transcriptomic and proteomic datasets from (28) and (29). Values represent the lowest and highest possible weighted average errors given the results of Flux Variability Analysis. Weighted average error values were calculated from flux maps generated using a scaling factor of 1.

| Dataset | Light Level | Weighted average error (%) |  |
| --- | --- | --- | --- |
|  |  | No gene expression weights | With gene expression weights |
| Mergner et al. Transcriptome | High | 169 – 180 | 14.7 – 17.1 |
|  | Low | 93.8 – 103 | 14.9 – 18.1 |
| Mergner et al. Proteome | High | 169 – 180 | 9.63 – 12.2 |
|  | Low | 93.8 – 103 | 8.75 – 10.9 |
| Klepikova et al. Transcriptome | High | 169 – 180 | 15.3 – 17.8 |

We wanted to confirm that these reductions in error are in fact dependent on weights calculated from gene expression data and not an artifact of the weighting procedure itself. Indeed, previous studies have used the application of randomized weights as a method of exploring different possible flux modes in a plant metabolic network (34). We found that substituting the leaf for the root proteomic dataset, and vice-versa, resulted in no reduction in weighted average error **(Supplemental Table 1)** compared to pFBA. Neither did randomization of the weight vector and subsequent optimization result in improvements in weighted average error, with the mean of the weighted average errors of 50 high-light condition flux maps generated with independent randomized weight vectors at a scaling factor of 1 being 201%, versus the unweighted error value of 169-180% for that condition.

### Increases in agreement between FBA-predicted and MFA-estimated fluxes are broadly distributed across central metabolism

Although there is variation among individual fluxes in the degree to which omic data integration improves agreement between predicted and experimentally derived values, the reduction in weighted error as a result of gene weight application is distributed broadly across the fluxes for which 13C-MFA estimates are available. If, for example, the improvement were due to a substantial decrease in one or a small number of high-flux reactions and a negligible decrease or even increase in error for other reactions **(Fig. 1)** the overall finding would be less striking and potentially less broadly applicable. The reductions in error are consistent not only across metabolic subsystems within a single FBA flux map, but also across alternative stoichiometric network structures. Initial pFBA-derived solutions for a model identical to that used to generate the other predictions except with unconstrained movement of protons show similar reductions in error **(Supplemental Table 2)**. Upon application of gene expression weights, this model converges to a similar value of weighted average error as other model configurations.

**Fig. 1.**
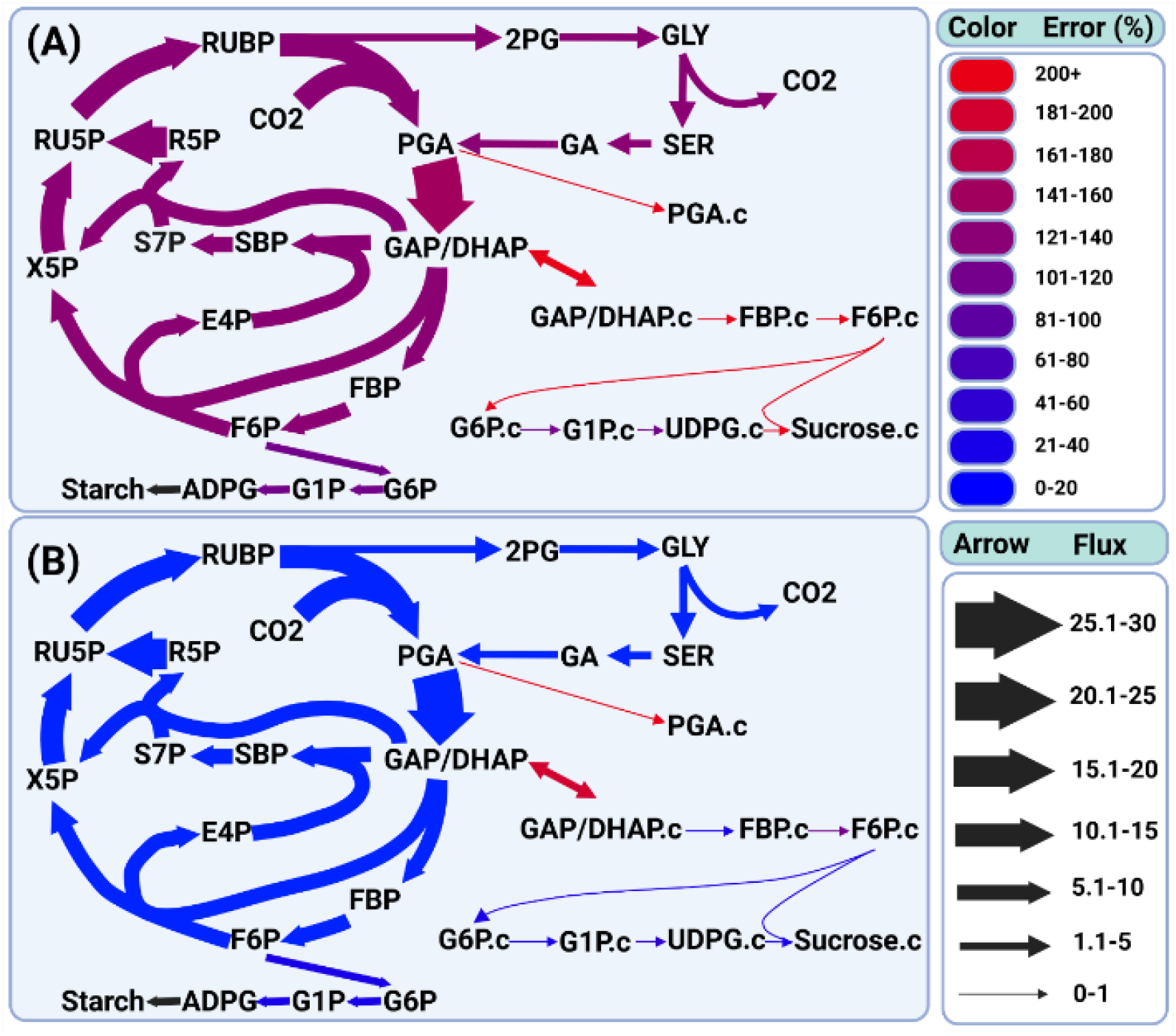
Percent errors relative to 13C-MFA derived fluxes of specific reactions in central metabolism before **(A)** and after **(B)** gene expression weight application. The error values in **(A)** are the **lowest** possible given FVA results and the values in **(B)** are the **highest** possible given FVA results. We see substantial decreases in errors associated with central carbon assimilation, as well as starch and sucrose synthesis. Since the 13C-MFA estimated fluxes from (26) do not feature the flux from ADPG to Starch, this flux lacks an estimated error and is therefore shown in black. Flux values are relative to the lowest flux in the network.

### Error reductions are a function of scaling factor parameter and are improved by the application of a tissue-specific median weight for reactions lacking Gene-Protein-Reaction terms

The magnitude of the gene expression weights calculated and applied by the present method depend on the magnitude of the scaling factor term, (see Methods, **Eq. 2**). The increased agreement between the FBA-predicted and MFA-estimated flux maps only manifests in the majority of cases for scaling factors of 0.05-0.1 or greater **(Fig 2)**. We also note that the relationship between the scaling factor value and the improved agreement is monotonic – that is, we do not see erratic increases and decreases as we increase the scaling factor value and, by extension, the strength of the assumed relationship between flux and gene expression. The necessity of a non-negligible scaling factor, the consistency of error improvement as the scaling factor is increased, and the similarity in the pattern of error improvement across multiple datasets as seen in **Fig 2** all suggest that real biological signal related to the partitioning of metabolic activity across the plant’s tissues is being extracted from the gene expression datasets. Finally, we observe that the flux maps generated using weight derived from the (28) proteomic dataset have noticeably better weighted average errors than flux maps generated using transcriptomic dataset **(Table 1; Fig. 2)**. This is consistent with the closer relationship between measured protein levels and metabolic fluxes than between transcripts and fluxes. It is also consistent with at least one other study’s attempts at integrating gene expression data into FBA in *E. coli* (24).

**Figure 2.**
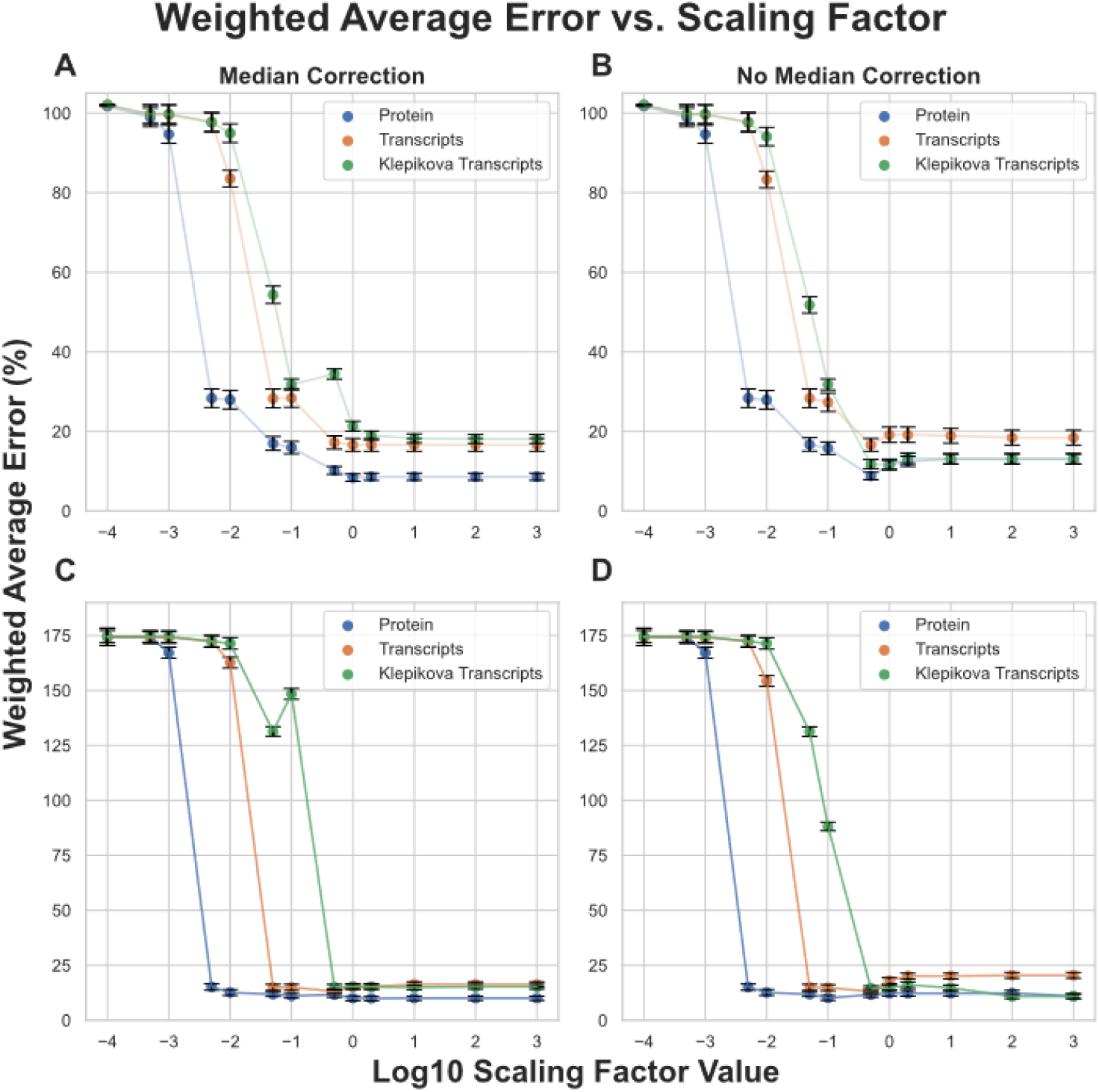
Weighted average errors of FBA predictions compared with MFA-estimated flux maps as a function of scaling factor value, light-level, and the presence or absence of a tissue-specific median weight correction. **(A)** Weighted average errors of flux maps generated using low-light constraints and with a tissue-specific median correction applied. **(B)** Weighted average errors of flux maps generated using low-light constraints and without a tissue-specific median correction applied. **(C)** Weighted average errors of flux maps generated using high-light constraints and with a tissue-specific median correction applied. **(D)** Weighted average errors of flux maps generated using high-light constraints and without a tissue-specific median correction applied. Upper and lower bars on each point represent the highest and lowest possible weighted average errors given FVA results, and the points themselves represent the average of these values.

In our initial formulation of the algorithm for generating gene expression weights, the weight of all reactions with no associated GPR was set to 1, since this is the implicit value of the coefficient for all reactions in a standard pFBA optimization. Since this runs the risk of introducing a systematic bias against using reactions that have associated GPRs, we attempted to counteract this effect by assigning all reactions lacking a GPR a weight corresponding to the median weight of all weighted reactions in the tissue in which those reactions are found. Comparing the results with and without the tissue-specific median weights for reactions without GPRs, we see slight improvements in the weighted average errors from a scaling factor of 1 onwards when using the transcriptomic and proteomic datasets from (28) **(Fig. 2)**, though the effect is not substantial, indicating that our method is robust to including or omitting the tissue-specific median weight correction.

### Changes in the carbon and energy economy upon application of gene expression weights

In addition to improving quantitative agreement between the FBA-predicted and MFA-estimated flux maps, the gene expression weighting procedure also generates flux maps that present a substantially different picture of carbon and energy metabolism in *Arabidopsis* leaves.

A consistent trend across high and low light FBA-predicted fluxes is a substantial decrease in leaf mitochondrial Electron Transport Chain (ETC) activity and overall flux in mitochondria-localized reactions in the light relative to nighttime ETC activity and overall flux (**Supplemental Table 3**). MFA and other recent work further supports low TCA cycle fluxes in photosynthesizing leaves (35–37). This decrease in mitochondrial activity goes hand-in-hand with a predicted decrease in the use of unusually high fluxes related to proline metabolism to indirectly support the consumption of excess reductant produced via the light reactions of photosynthesis. Alongside this decrease in mitochondrial activity is a decrease in the ratio of cyclic electron flow (CEF) to linear electron flow (LEF) in the chloroplast **(Table 2)**. Although reliable empirical measurements of this CEF/LEF ratio are difficult to obtain, previous studies have shown that a C3 plant like *Arabidopsis* relying on cyclic electron flow to bring the ratio of ATP/NADPH produced up to that needed for normal growth would have a CEF amounting to ∼13% of LEF (38). Due to the presence of other balancing mechanisms, such as the malate valve (39), this 13% value would represent an upper bound on stoichiometrically predicted values for CEF/LEF. Application of gene expression data see decreases the CEF/LEF ratios in all but one FBA-predicted flux map to values much closer to the expected ∼13% upper bound than are predicted using conventional pFBA **(Table 2)**.

**Table 2.**
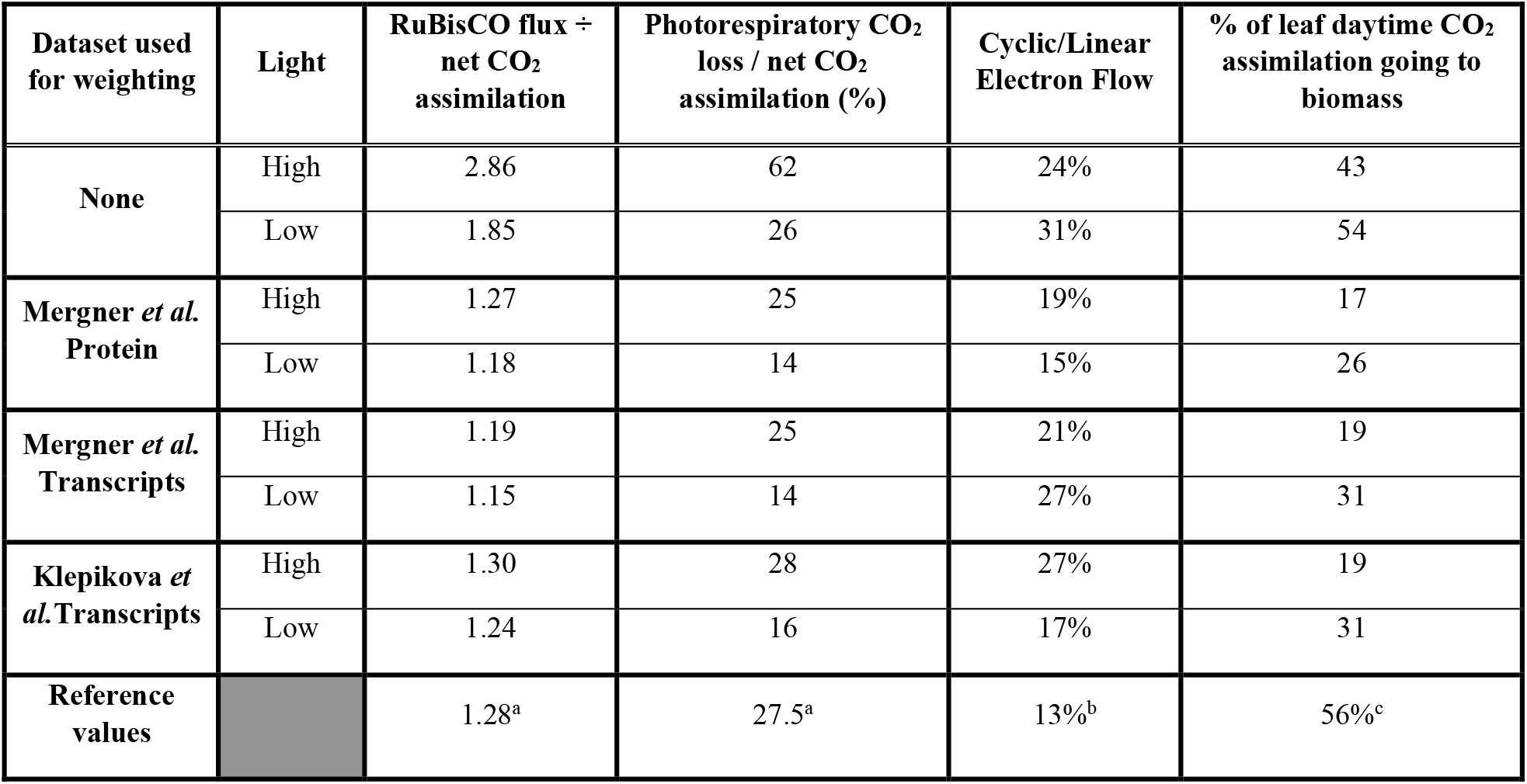
Several measures of carbon and energy utilization derived from the predicted flux maps with and without gene expression weighting applied. Reference values: a, (26); b, (38); c, (40).

| Dataset used for weighting | Light | RuBisCO flux ÷ net CO<sub>2</sub> assimilation | Photorespiratory CO<sub>2</sub> loss / net CO<sub>2</sub> assimilation (%) | Cyclic/Linear Electron Flow | % of leaf daytime CO<sub>2</sub> assimilation going to biomass |
| --- | --- | --- | --- | --- | --- |
| None | High | 2.86 | 62 | 24% | 43 |
|  | Low | 1.85 | 26 | 31% | 54 |
| Mergner *et al.* Protein | High | 1.27 | 25 | 19% | 17 |
|  | Low | 1.18 | 14 | 15% | 26 |
| Mergner *et al.* Transcripts | High | 1.19 | 25 | 21% | 19 |
|  | Low | 1.15 | 14 | 27% | 31 |
| Klepikova *et al.* Transcripts | High | 1.30 | 28 | 27% | 19 |
|  | Low | 1.24 | 16 | 17% | 31 |
| Reference values |  | 1.28<sup>a</sup> | 27.5<sup>a</sup> | 13%<sup>b</sup> | 56%<sup>c</sup> |

From (26) we have MFA-derived estimates of %vpr, or the rate of photorespiratory CO_2_ release via glyoxylate decarboxylation, as well as the ratio of RuBisCO carboxylation flux to net CO_2_ assimilation in the leaf. The unweighted flux predictions for the high and low light conditions disagree substantially with these estimates **(Table 2)**. However, application of gene expression weights consistently brings estimates of these parameters into close agreement with MFA-derived values. The integration of gene expression also changes the predicted efficiency with which *Arabidopsis* converts atmospheric CO_2_ into biomass **(Table 2)**. For comparison with these predicted efficiencies, we used the empirical *A. thaliana* biomass, leaf area, and gas exchange data reported in (40) to calculate that approximately 56% of the net CO2 assimilation in illuminated leaves ends up in incorporated into biomass, which is closer to the value in our unweighted flux predictions than our weighted flux predictions, although it should be noted that these data were gathered from a hydroponic system.

## Discussion

13C-MFA is broadly accepted as being the most reliable method for estimating metabolic flux maps *in vivo* due to its ability to make use of substantial amounts of isotopic labeling data to arrive at well-supported flux maps in small-to medium-scale networks (5). However, the technique’s utility is limited by the substantial experimental effort that goes into the generation of each individual flux map. FBA, with its requirement of much less experimental data, has become the method of choice for more exploratory or predictive metabolic modeling studies. The implicit assumption is usually that the predictions of FBA – or at least the range of its predictions in cases where a unique solution is not provided – agree with those we would arrive at if we were able to conduct a 13C-MFA study. This makes our optimization procedures when performing FBA and validation of FBA models against MFA results of vital importance. The method presented here, by bringing FBA-predicted fluxes into line with MFA-estimates represents a step in the direction of higher-confidence FBA flux maps.

One limitation, as well as motivation, for the present study is the lack of a large set of 13C-MFA datasets in plants and other multi-tissue eukaryotic systems. Systems like *E. coli* have multi-omic datasets consisting of transcriptomic, proteomic, and fluxomic measurements (16) that have been utilized to empirically infer the relationship between gene expression and metabolic fluxes. This empirical training can then be used to more accurately predict fluxes in new contexts (24). The sparsity of 13C-MFA data in more complex systems makes such an approach currently impossible.

An interesting theoretical aspect of the present approach is its simplicity, the only variable parameter being a single scaling factor that controls the magnitude of the penalty weights. That the assumption of a consistent value relating the relative abundances of transcripts or proteins in different tissues to the “preference” of an organism to partition flux among particular reactions can result in substantial improvement in error was of great interest. This observation is interesting to consider in light of the complexity of the relationship between measures of gene expression – transcriptomic and proteomic abundances – and flux. Particularly when making biotechnological interventions in a system to modify its metabolism, there is often an assumed strong linear relationship between transcription, translation, and, ultimately, metabolic flux, but the reality is rarely so simple. Although moderate correlations between transcript and protein abundances have been demonstrated across many systems, the degree of correlation varies across systems and experimental contexts (41, 42). The correlation between these datatypes and rates of central metabolic reactions, which carry the large majority of total metabolic flux, is weaker still (43). Some previous studies found that changes in the gene expression related to individual reactions typically do not correlate well with changes in fluxes (24, 44), with some central metabolic fluxes in particular showing a negative correlation between gene expression change and flux change. In both cases, gene expression data related to reactions were compared within the same cell type or tissue; in our study, we avoid this comparison, comparing instead inter-tissue abundances, mirroring the long-standing practice in the literature of inferring relative metabolic activity in different tissues by their transcript and protein investment in relevant pathway steps. It may be the case that it is only by considering gene expression on an inter-tissue basis in the context of the entire complex stoichiometric network underlying metabolism that predictive gains from including gene expression evidence can be properly realized.

Future work should aim to expand the number of available datasets, and the experimental conditions and genotypes for which they are gathered, in order to enable more thorough evaluation of methods like the one presented in this paper. Building on the work of Ma *et al*. (26), experimental improvements and refinements of the underlying network architecture of central carbon metabolism have been introduced in the context of 13C-MFA in *Camelina sativa* (35, 36) and *Nicotiana tabacum* (45). In the present study the Ma et al. 2014 flux maps are used without change and we adopted a highly curated *A. thaliana* genome-scale model from which to construct the whole-plant model. This approach precluded the possibility of our reanalyzing the MFA-estimated flux map, or constructing a new purpose-built genome-scale model, making the MFA-to-FBA comparison more favorable. However, in future studies a combination of MFA network refinements, expanded datasets, and further improvements in the flux estimation procedures holds promise for improving the fidelity of the 13C-MFA comparison data. On the FBA side, the use of more detailed growth and composition measurements for FBA along with more detailed representation of different tissue types will potentially allow for more biologically accurate and representative FBA flux map predictions. These improvements in both MFA-estimation and FBA-prediction of flux maps, along with an expansion in the number of available 13C-MFA datasets against which to compare FBA predictions, will allow for more extensive validation of the method described in this paper as well as other methods aiming to incorporate omic datasets into flux prediction.

A distinct aspect of the proposed method is its demonstrated ability to bring FBA-predicted fluxes in line with MFA-estimated fluxes across multiple input datasets, model architectures, and using multiple independent gene expression datasets. Our hope is that methods for incorporating transcriptomic and proteomic data may advance this field to the point where FBA-predicted flux maps can be used with high-confidence for practical engineering goals. This, combined with the automated reconstruction of GEMs from genomic and biochemical databases (46) suggests a future with rapid turnaround from the initial identification of an organism of interest to metabolic flux predictions and rational genetic engineering to achieve biotechnological aims.

## Acknowledgements

We would like to thank Dr. Doug Allen for permission to adapt **Figure 3** from (26) for use in **Figure 1** in this publication. Figures 1 and 3 were created with BioRender.com. This research was supported by the Office of Science (BER), U.S. Department of Energy, Grant no DE-SC0018269 (J.A.M.K. and Y.S-H.). This work is supported, in part, by the NSF Research Traineeship Program (Grant DGE-1828149) to J.A.M.K. This publication was also made possible by a predoctoral training award to J.A.M.K. from Grant T32-GM110523 from National Institute of General Medical Sciences (NIGMS) of the NIH. Its contents are solely the responsibility of the authors and do not necessarily represent the official views of the NIGMS or NIH.

## Conflict of interest statement

The authors have no conflicts of interest to declare.

## Notes

https://github.com/Gibberella/ArabidopsisGeneExpressionWeights

### Competing Interest Statement

The authors have declared no competing interest.

