## Supplemental Information for "Accurate flux predictions using tissue-specific gene expression in plant metabolic modeling"

**Supplementary Information for “Accurate flux predictions using tissue-specific gene expression in plant metabolic modeling”**

Joshua A.M. Kaste, Yair Shachar-Hill

Corresponding author: Yair Shachar-Hill

**Supplementary Methods**

**Datasets and Omic Data Processing**

Sample IDs and SRR numbers for all transcriptomic and proteomic datasets used in this study can be found in **Supplemental Table 4**.

The raw RNA-seq data for all 137 samples in (1) was trimmed using the *fastp* algorithm (2) and then aligned to the TAIR10 Arabidopsis thaliana genome obtained from ensembl plants using the *salmon* algorithm (3–5). RNA-seq reads from (6) were taken directly from the published supplemental material. Library normalization was performed on the RNA-seq datasets from both (1, 6) using the DeSeq2 procedure (7) and averages of the normalized transcript abundance values across replicates from (1) were used. Intensity-based Absolute Quantification (IBAQ) (8) values for the proteomic data from (6) were divided by the sum total intensity across all protein signals measured for a given sample to normalize them.

**Error Evaluation and Statistical Analysis**

To evaluate whether the error values for measured reactions in individual flux maps generated using gene expression weights were statistically significantly different from the errors without application of these weights, the Wilcoxon signed-rank test was used (10). The Bonferroni-Holm multiple testing correction (11) was used to correct the family-wise α of all hypothesis tests to 0.01, where each hypothesis test is asking, by the Wilcoxon signed-rank test, whether the differences between a given FBA-predicted flux map (e.g. the flux map generated using protein-derived gene expression weights and a Scaling Factor of 1) and our MFA-estimated flux map could be attributed to random chance. In the high light condition 13C-MFA flux map from (9), the flux through the malate dehydrogenase reaction was reported as exactly 0 – as this made its error undefined, it was excluded from the high light condition’s error calculation.

**Flux Variability Analysis**

Flux Variability Analysis (FVA) (12) was used to determine the maximum and minimum values possible for each of the fluxes included in the error calculation, subject to the following constraint:

$$\begin{aligned} c*v=opt \#\left( 1 \right) \end{aligned}$$

Where *c* is the vector of all weight coefficients, *v* is the vector of all fluxes, and *opt* is the value of the objective function determined by the initial optimization procedure. As shown in **Supplemental Dataset 3,** the some of the fluxes included in the error function are not uniquely defined, such that they can vary up and down without violating **Eq. 1**. To account for this variation, maximum and minimum weighted average errors were calculated, where the minimum and maximum errors correspond to the smallest and largest weighted average errors possible for a flux map given the maximum/minimum values for all evaluated fluxes. FVA was performed in MATLAB using the COBRA Toolbox (13) and Gurobi™ 8.1.1 (14).

**Model Constraints**

Light uptake and photosynthetic activity were restricted to the leaf tissue and mineral uptake was restricted to root tissue. Inter-tissue transport and day/night continuity of metabolites were defined and constrained as in Cheung & Shaw 2018 (15) as were ATP and NADPH maintenance flux values. Biomass compositions of leaf, stem, and root were taken from Dal’Molin 2015 (16), based on (17). Reactions were added to produce biomass components that appear in the Dal’Molin biomass equations (16) but not in the core metabolic model (18); this involved adding subnetworks of missing reactions for several components and single summary reactions for others. Cytosolic pentose phosphate pathway reactions were also added to the model. All reactions were converted to irreversible form, wherein all reversible reactions were converted to independent forward and reverse reactions, prior to solving. This is simply to ensure that all fluxes, including those representing the reverse flux of a reversible reaction, take values that are zero or positive.

In order to generate predictions corresponding to the high-light and low-light flux maps reported in (9), the vo/vc, or ratio of ribulose 1,5-bisphosphate carboxylase/oxygenase (RuBisCO) oxygenation activity to its carboxylation activity, and the ratio of starch to sucrose synthesis were both constrained to the values estimated in that study.

**Supplementary Tables**

Table S1. Reductions in weighted average error with application of gene expression weights derived from gene expression data with incorrect tissue specification or randomized values

| Dataset | Light Level | Weighted Average Error (%) | |
| --- | --- | --- | --- |
|  |  | **Without Gene Expression Weights** | **With Leaf/Root Flipped Gene Expression Weights** |
| **Mergner et al. Transcriptome** | High | 168 – 180 | 182 - 215% |
|  | Low | 93.8 – 103 | 155 - 183 |
| **Mergner et al. Proteome** | High | 168 – 180 | 249 - 295 |
|  | Low | 93.8 – 103 | 132 - 159 |
| **Klepikova et al. Transcriptome** | High | 168 – 180 | 87.9 - 109 |
|  | Low | 93.8 – 103 | 97.2 – 119 |

^a^Weighted average errors are calculated from flux maps generated using a scaling factor of 1.

**Table S2.** Reductions in weighted average errors with an alternate model architecture

| Dataset | Light Level | Weighted Average Error (%) | |
| --- | --- | --- | --- |
|  |  | **Without Gene Expression Weights (%)** | **With Gene Expression Weights (%)^a^** |
| **Mergner et al. Transcriptome** | High | 127 - 135 | 11.8 – 14.2 |
|  | Low | 66.1 - 73.8 | 17.3 – 20.0 |
| **Mergner et al. Proteome** | High | 127 - 135 | 10.6 - 12.8 |
|  | Low | 66.1 - 73.8 | 9.00 - 11.2 |
| **Klepikova et al. Transcriptome** | High | 127 - 135 | 13.9 - 16.5 |
|  | Low | 66.1 - 73.8 | 21.0 - 23.4 |

^a^Weighted average errors are calculated from flux maps generated using a scaling factor of 1.

**Table S3.** Ratios of Day vs. night leaf mitochondrial fluxes and Electron Transport Chain fluxes in flux maps with and without integration of gene expression evidence.

| Dataset | Light Level | Ratio of total mitochondrial flux in day vs. night | | Ratio of mitochondrial ATP synthase flux in day vs. night | |
| --- | --- | --- | --- | --- | --- |
|  |  | **Without Gene Expression Weights** | **With Gene Expression Weights^a^** | **Without Gene Expression Weights** | **With Gene Expression Weights^s^** |
| **Mergner et al. Transcriptome** | High | 1.08 | 0.169 | 1.15 | 0.0888 |
|  | Low | 1.20 | 0.287 | 1.20 | 1.94 * 10^-5 |
| **Mergner et al. Proteome** | High | 1.08 | 0.183 | 1.15 | 6.17 * 10^-5 |
|  | Low | 0.872 | 0.483 | 1.20 | 1.87 * 10^-5 |
| **Klepikova et al. Transcriptome** | High | 1.08 | 0.148 | 1.15 | 0.122 |
|  | Low | 0.872 | 0.391 | 1.20 | 0.537 |

^a^Values for weighted cases calculated from flux maps generated using a scaling factor of 1.

Table S4. Sample IDs and SRR numbers used in the present study.

| **Dataset** | **Reference** | **Leaf** | **Stem** | **Root** |
| --- | --- | --- | --- | --- |
| **Mergner et al.** | (6) | AP13,  AP14 | AP11 | AP17 |
| **Klepikova et al.** | (1) | SRR3581680,  SRR3581846 | SRR3581705,  SRR3581871 | SRR3581356,  SRR3581836 |

**Supplementary Datasets Descriptions**

**Dataset S1 (separate file).** Mapping between reactions in FBA model and Ma *et al.* MFA flux map (9) reactions.

**Dataset S2 (separate file).** Full model used for FBA in .xls format.

**Dataset S3 (separate file).** Flux Variability Analysis results.

**Dataset S4 (separate file).** Flux vectors generated for this study.

**Dataset S5 (separate file).** Weighted average error calculations done for this study.

**Dataset S6 (separate file).** Penalty weights used in this study.
